# A chemical probe to modulate human GID4 Pro/N-degron interactions

**DOI:** 10.1101/2023.01.17.524225

**Authors:** Dominic D.G Owens, Matthew E.R Maitland, Aliakbar Khalili Yazdi, Xiaosheng Song, Martin P. Schwalm, Raquel A.C Machado, Nicolas Bauer, Xu Wang, Magdalena M. Szewczyk, Cheng Dong, Aiping Dong, Peter Loppnau, Matthew F. Calabrese, Matthew S. Dowling, Jisun Lee, Justin I. Montgomery, Thomas N. O’Connell, Chakrapani Subramanyam, Feng Wang, Matthieu Schapira, Stefan Knapp, Masoud Vedadi, Jinrong Min, Gilles A. Lajoie, Dalia Barsyte-Lovejoy, Dafydd R. Owen, Caroline Schild-Poulter, Cheryl H. Arrowsmith

## Abstract

The CTLH complex is a multi-subunit ubiquitin ligase complex that recognizes substrates with Pro/N-degrons via the substrate receptor GID4. Recently, focus has turned to this complex as a potential mediator of targeted protein degradation, but the role GID4-mediated substrate ubiquitylation and proteasomal degradation plays in humans has thus far remained unclear. Here, we report PFI-7, a potent, selective, and cell-active chemical probe that antagonizes Pro/N-degron binding to human GID4. Use of PFI-7 in proximity-dependent biotinylation enabled the identification of dozens of endogenous GID4-interacting proteins that bind via the GID4 substrate binding pocket, only a subset of which possess canonical Pro/N-degron sequences. GID4 interactors are enriched for nuclear and nucleolar proteins including RNA helicases. GID4 antagonism by PFI-7 altered protein levels of several proteins including RNA helicases as measured by label-free quantitative proteomics, defining proteins that are regulated by GID4 and the CTLH complex in humans. Interactions with GID4 via Pro/N-degron pathway did not result in proteasomal degradation, demonstrating that CTLH interactors are regulated through a combination of degradative and non-degradative functions. The lack of degradation of GID4 interactors highlights potential challenges in utilizing GID4-recruiting bifunctional molecules for targeted protein degradation. Going forward, PFI-7 will be a valuable research tool for defining CTLH complex biology and honing targeted protein degradation strategies.

## Introduction

The half-life or stability of most intracellular proteins is governed by the presence of short linear sequences known as degrons^1–4^. Degrons at the N- or C-terminus of a protein, known as N- and C-degrons, or internal degrons such as PEST sequences, recruit distinct E3 ubiquitin ligases to polyubiquitylate substrates and target them for proteasomal degradation^1,5^. E3 ligase-mediated polyubiquitylation of proteins via ubiquitin Lysine (K)-48 chains or K48-K11 branched chains typically leads to substrates being targeted to the 26S proteasome for subsequent degradation^6^. Degrons may be recognized by E3 ligases directly or may undergo processing such as proteolytic trimming by amino peptidases, addition of terminal amino acids, or other posttranslational modifications including phosphorylation^1–4^. Degradation of proteins with distinct degrons occurs at different rates depending on the specific downstream enzymatic machinery recruited by each degron^2,7^. Collectively, degron sequences fine-tune the abundance of intracellular proteins and further insights into degron recognition pathways will likely inform therapeutic efforts that employ targeted protein degradation^8^.

The Pro/N-degron pathway is a recently identified pathway that recognizes N-terminal degrons containing unmodified Proline residues (Pro/N-degrons), either at position one or two of the sequence^9,10^. The Pro/N-degron pathway in *Saccharomyces cerevisiae* allows cells to quickly adapt their metabolism and maintain homeostasis^9^. Pro/N-degron-containing gluconeogenic enzymes are recognized by the Gid4 β-barrel substrate binding pocket and subsequently ubiquitylated by a multi-subunit E3 ligase complex called the Glucose-induced degradation (GID) complex, targeting them for proteasomal degradation^9^. Many of the orthologous GID complex subunits exhibit conserved sequence homology and function between yeast and humans, but as several subunits contain a C-terminal to LisH (CTLH) domain, the complex is commonly referred to as the CTLH complex in humans (Fig. 1A)^11^. Notable differences between the yeast and human complexes have been observed^12^. In *S. cerevisiae*, Gid4 is interchangeable with other substrate receptors including Gid10 and Gid11^13–15^, while GID4 is thus far the only CTLH complex substrate receptor identified in humans, despite some substrates being degraded in a GID4 independent manner^16^. In humans, the CTLH complex is implicated in a wide variety of cellular processes including autophagy, development, cell cycle regulation, and primary cilium function^12^. Mutations in, and over expression of CTLH complex members are also common in some cancers^17^. Mutations in the substrate binding pocket of GID4 that are likely to disrupt substrate binding have been reported in glioma, breast and pancreatic carcinoma samples^10^. Together, this suggests that the role of the CTLH complex in humans has expanded beyond the initial function evolved in yeast, possibly through acquisition of a wider set of ubiquitylation substrates.

**Figure 1.**
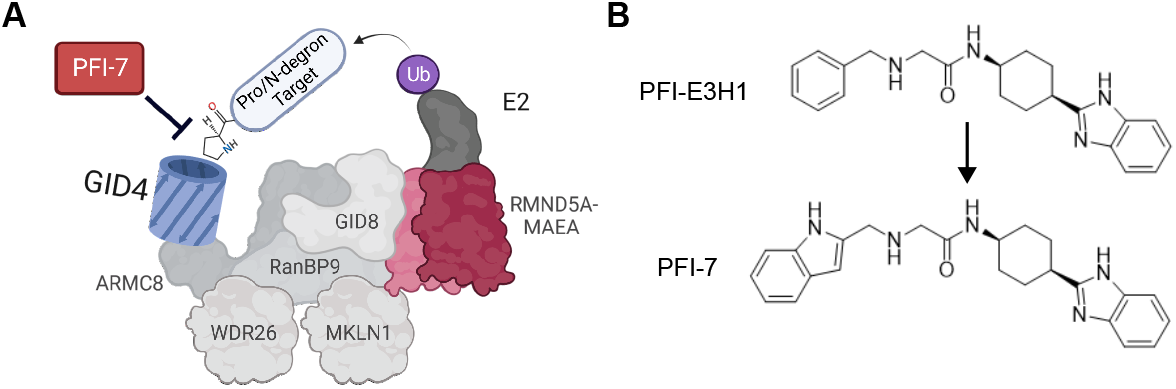
A) Schematic of GID4 bound to the human CTLH complex with E2 enzyme and ubiquitin shown. The recognition of degron-containing substrates by GID4 is inhibited by GID4 antagonist PFI-7. B) Chemical structures of PFI-E3H1 (GID4 E3 handle) and PFI-7 (GID4 chemical probe).

Substrates of the CTLH complex have been more challenging to identify in humans than in yeast. The first human CTLH substrate to be identified was HMG box protein 1 (HBP1)^18^, a transcription factor responsible for inhibiting expression of pro-proliferative cell cycle regulators. HBP1 is ubiquitylated and degraded independently of GID4^16,18^, suggesting degradation is mediated via an alternative recognition subunit in CTLH other than GID4. Since then, a handful more degradative substrates of the complex have been suggested including BICC1^19^, MKLN1^20^, LMNB2^21^, RAF1^22^, and ZMYND19^16^. Intriguingly, ubiquitylation of glycolysis enzymes PKM2 and LDHA is dependent on the CTLH complex^23^, indicating a conserved role for GID/CTLH complex in regulating metabolism in yeast and humans. However, ubiquitylation of PKM2 and LDHA does not result in their degradation, instead it reduces their activity to decrease glycolytic metabolism^23^. None of the human CTLH complex substrates characterized so far contain either a canonical Pro/N-degron, or the more flexible degron sequences that were recently determined to also be recognized by GID4 in vitro^24,25^. Bioinformatic prediction has been used to identify potential substrates based on the presence of a canonical Pro/N-degron consensus sequence^15^, and some of these putative substrates have been shown to interact with CTLH complex^21^; however, a global assessment of GID4-dependent interactors of the CTLH complex and the role GID4 plays in proteome regulation in humans is so far lacking.

To address this knowledge deficit, we developed a selective antagonist of human GID4. PFI-7 is a chemical probe that binds within the β-barrel of GID4 substrate binding pocket, disrupting its interaction with the canonical Pro/N-degron peptide in cells. Using proximity-dependent biotinylation (BioID2-GID4) coupled with liquid chromatography tandem mass spectrometry (LC-MS/MS), we identify dozens of endogenous GID4 interactors whose interaction with GID4 is reduced by PFI-7. GID4-dependent interactors are enriched for nucleolar proteins and proteins associated with RNA metabolism including RNA helicases. A subset of interactors exhibit canonical Pro/N-degrons, while the majority do not, implicating versatility in substrate recruitment to the CTLH complex by GID4. GID4 regulated abundances of proteins including RNA helicases, while levels of several GID4 interactors did not change after GID4 inhibition, showing that recruitment to GID4 via Pro/N-degron sequences is not sufficient to promote proteasomal degradation. This proteome-wide assessment sheds new light on GID4-mediated CTLH complex interactors and identifies proteins regulated by GID4/CTLH. Furthermore, this work positions PFI-7 as an important research tool to further dissect mechanistic details and biological roles of the CTLH complex in health and disease. Our data also provide insights into proteome regulation by CTLH/GID4 that will inform future efforts to establish GID4 recruitment reagents or drugs for targeted protein degradation^8,26^.

## Results

### Discovery of a potent ligand targeting GID4 substrate binding pocket

In order to discover a chemical probe to explore the E3 ligase activity of the CTLH complex, we targeted its evolutionarily conserved substrate receptor, GID4 (Fig. 1A). Affinity selection mass spectrometry and subsequent medicinal chemistry optimization led to the discovery of the moderately potent ‘E3 Handle’ compound, PFI-E3H1 (Khalili Yazdi et al., In preparation, Fig. 1B). After slight modification to the PFI-E3H1 scaffold to improve anticipated potency, the compound PFI-7 was selected for further characterization. A co-crystal structure shows PFI-7 bound deep within the peptide binding pocket that recognizes Pro/N-degrons^10^ (Fig. 2A, Supp. Table 1). PFI-7 binds to GID4 with a Kd of 79 ± 7 nM, as determined by surface plasmon resonance (SPR, Fig. 2B). Furthermore, PFI-7 can directly displace the Pro/N-degron peptide PGLWKS from the GID4 binding pocket in a peptide displacement assay (Fig. 2C, Kdisp 4.1 ± 0.2 μM). Consistent with these data, PFI-7 binds through a network of hydrophobic and H-bond interactions (Fig. 2A), with the benzimidazole moiety surface exposed, indicating a site on PFI-7 that will likely tolerate derivatization. Together this biophysical characterization reveals that PFI-7 binds to the GID4 substrate binding pocket and disrupts Pro/N-degron binding.

**Figure 2.**
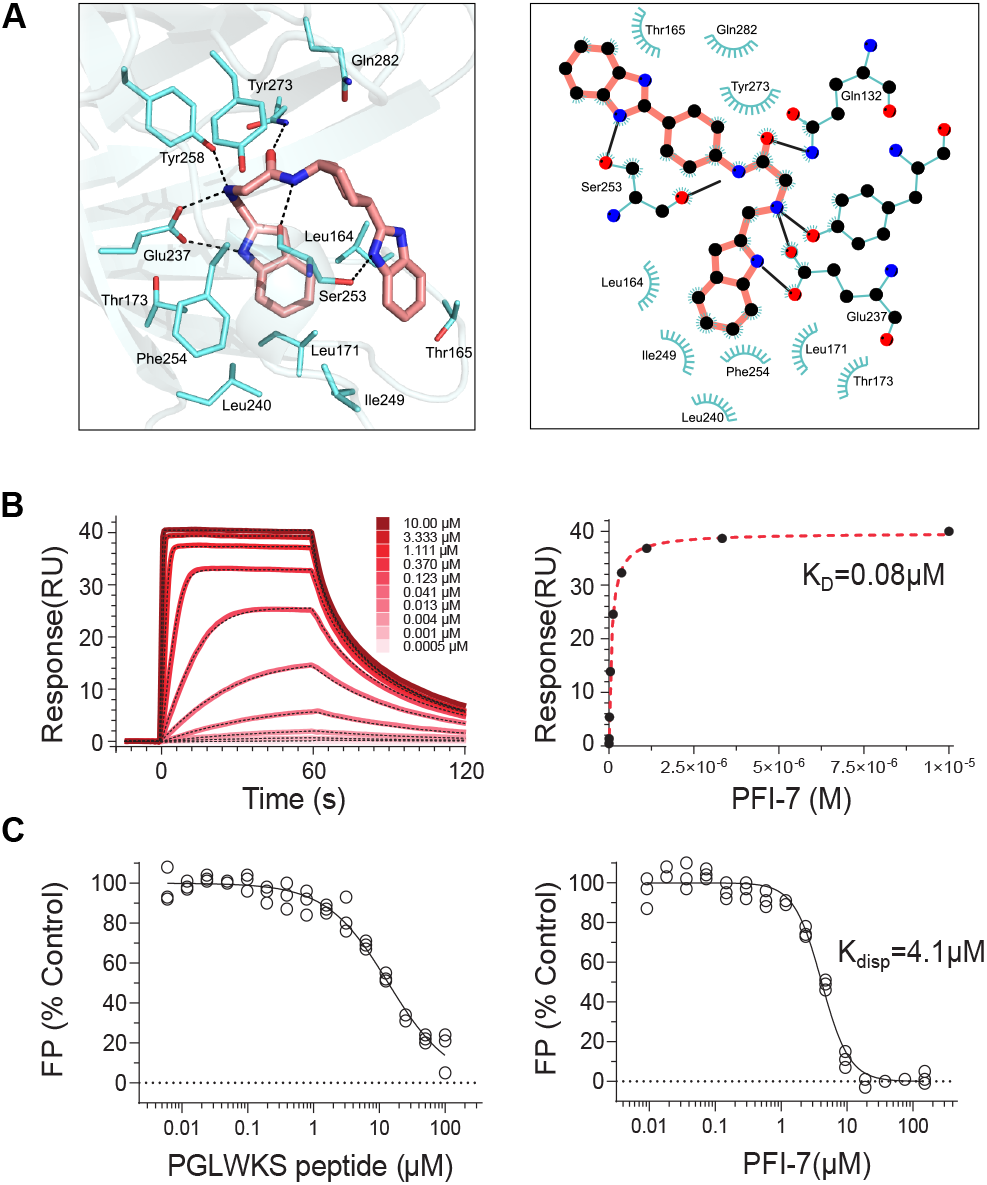
A) Crystal structure (left panel) of PFI-7 (shown in pink) bound to GID4 (shown in grey and cyan). Right panel shows in two dimensions the key interactions between GID4 and PFI-7. B) SPR analysis of the binding of PFI-7 to GID4. Serially diluted compounds were flown over immobilized GID4 and the resulting response units (RU) signals were monitored. Left, a representative sensorgram for each compound is shown (solid red lines) with the corresponding 1:1 binding kinetics fit indicated (dashed black lines). Right, the steady state response obtained from the data (left) is indicated (black circles) along with the corresponding steady state 1:1 binding model fitting (dashed red-lines). Experiments were performed in triplicate (n=3). C) FP-based peptide displacement assay. Compounds were tested for competing with the fluorescein-labeled PGLWKS peptide for binding to GID4. Left, unlabelled PGLWKS peptide was used as a control (0.006 to 100 μM). Right, PFI-7 was used at concentrations ranging from 0.01 to 150 μM. All experiments were performed in triplicate (n=3) and the calculated Kdisp value is indicated.

### PFI-7 is a potent GID4 antagonist in cells

To profile target engagement in a cellular context and quantify the extent to which PFI-7 engages and antagonizes GID4 in live cells, we employed NanoLuciferase (NanoLuc) bioluminescence resonance energy transfer (NanoBRET) protein-protein interaction (PPI) assays. In this assay, NanoLuc protein that acts as an energy donor was N-terminally tagged with PGLWKS peptide to mimic an endogenous Pro/N-degron (Fig 3A). GID4 was C-terminally tagged with haloalkane dehalogenase (HaloTag) that irreversibly binds to a cell-permeable chloroalkane-modified 618nm fluorophore, the BRET energy acceptor. A dose-dependent decrease in GID4-HaloTag binding to PGLKWS-NanoLuc was observed upon treatment of HEK293T cells with PFI-7 (IC50 = 0.57 ± 0.04 μM) (Fig. 3B), indicating that PFI-7 inhibits Pro/N-degron binding by GID4 in live cells. In a complementary assay configuration, we developed a tracer compound consisting of a derivatized PFI-E3H1 conjugated to a 618nm fluorophore, (GID4-Tracer) that dose-dependently bound to NanoLuc-tagged GID4 in live cells (Fig. 3C-E). PFI-7 effectively competed with the GID4-NanoLuc and GID4-Tracer interaction in a dose dependent manner (GID4-N IC50 = 0.35 ± 0.07 μM, GID4-C IC50 = 0.28 ± 0.05 μM) (Fig. 3F). There were minimal cytotoxic effects following treatment of a panel of cell lines with PFI-7 or PFI-E3H1 up to 10 μM over 3 days (Extended data Fig 1). Together, these data show that PFI-7 engages GID4 in live cells to inhibit Pro/N-degron binding and is a suitable tool compound for use in cellular studies to interrogate GID4-mediated recruitment to the CTLH complex.

**Figure 3.**
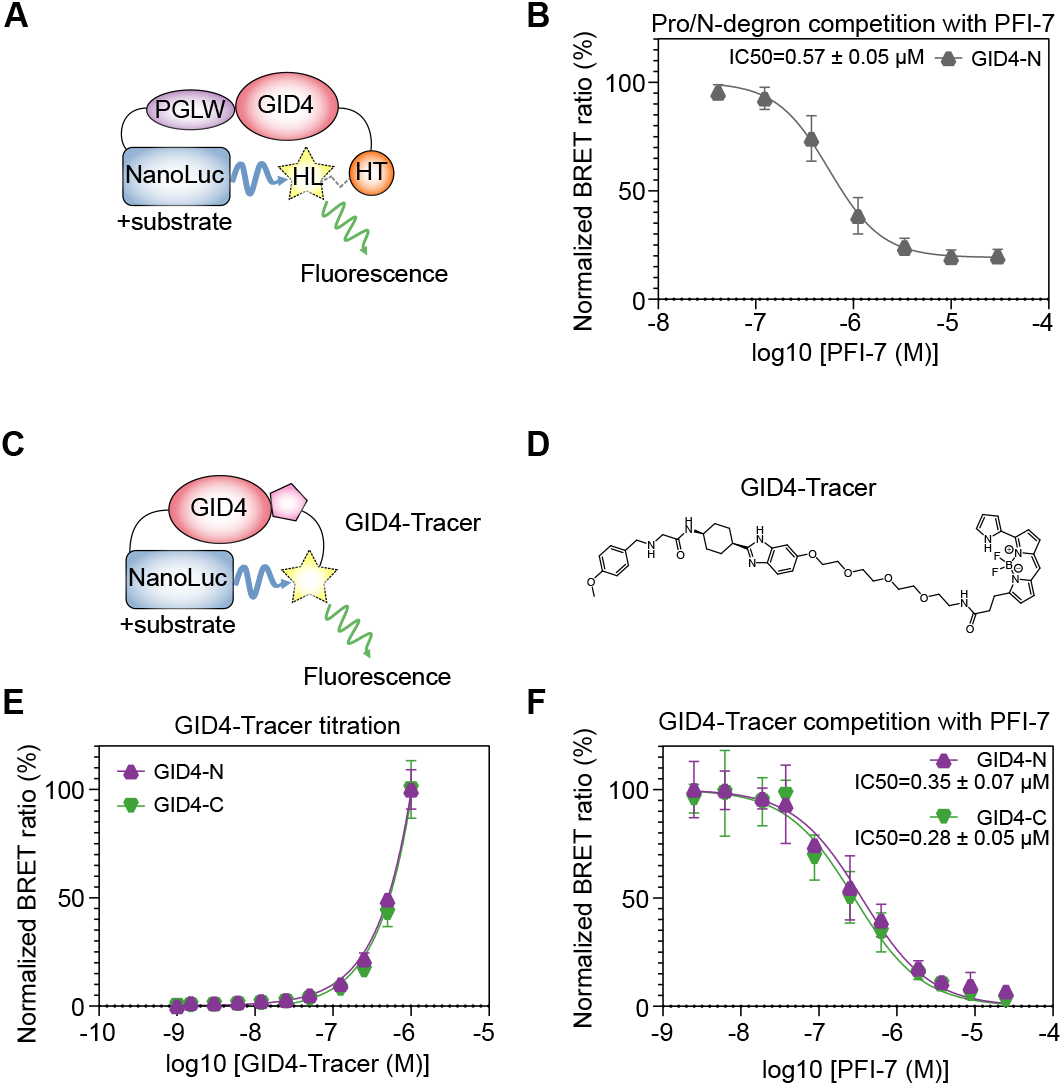
A) NanoBRET protein-protein interaction assay to quantify inhibition of GID4 Pro/N-degron binding by PFI-7. Schematic of PGLWKS degron-tagged NanoLuciferase (NanoLuc, donor) and C-terminally tagged GID4-haloalkane dehalogenase (HaloTag, HT) irreversibly bound to a cell-permeable 618nm fluorochrome (HaloLigand, HL, acceptor). B) BRET ratio as a percentage of vehicle (DMSO) control in cells treated with increasing concentrations of PFI-7. Data are from three independent experiments (n=3). C) NanoBRET target engagement assay utilizing GID4-Tracer compound. Left, schematic of GID4-NanoLuc fusion protein (donor) and a GID4-Tracer compound consisting of a GID4-binding moiety related to PFI-E3H1 derivatized with a 618nm fluorophore (acceptor). D) Right, chemical structure of GID4-Tracer compound. E) Titration of GID4-Tracer compound binding to N or C-terminally NanoLuc tagged GID4. BRET ratio is shown as a percentage of the highest measured value. F) Competition of GID4-Tracer compound binding to N or C-terminally NanoLuc tagged GID4 using GID4 antagonist PFI-7. BRET ratio is shown as a percentage of vehicle (DMSO) treated cells.

### GID4 interacts with proteins associated with RNA binding, transcription, splicing, and chromatin

The interactome of GID4 has never been determined using GID4 as a bait. Thus, to elucidate the GID4 interactome, proximity-dependent biotinylation was performed followed by protein identification by liquid chromatography tandem mass spectrometry (LC-MS/MS) (Fig. 4A). The GID4 fusion protein (HA-myc-BioID2-GID4, hereafter referred to as BioID2-GID4) preserves the GID4 C-terminal anchor required for CTLH complex binding via ARMC8a^14^ (Extended data Fig. 2A). BioID2-GID4 was functional as it interacted with the endogenous CTLH complex by coimmunoprecipitation (Extended data Fig. 2B). Upon doxycycline induction, BioID2-GID4 elicited biotinylation of proteins in an exogenous biotin-dependent manner (Extended data Fig. 2C). BioID2-GID4 showed similar localization to HA-GID4 as measured by immunofluorescence and western blot (Extended data Fig. 2D, E, and ref [20]). To interrogate the transient GID4 interactors that are degraded upon ubiquitylation by the CTLH complex, the GID4 interactome was analyzed in the presence and absence of proteasome inhibitor MG132^27^. Samples segregated based on GID4 expression in uniform manifold approximation and projection (UMAP) analysis, while proteasome inhibitor contributed little to sample clustering (Extended data Fig. 2F). Using a high confidence threshold, 196 proteins were significantly enriched in BioID2-GID4 samples compared to BioID2 alone (Saint probability [SP] ≥ 0.9, Fig. 4C, Supp. Table 2). Two members of the CTLH complex were identified as GID4 interactors, RANBP9 and MKLN1 (Fig. 4B, C). Twenty of the high-confidence GID4 interactors have previously been described as interactors of the CTLH complex in the BioGRID database^28^ including DDX50 (Extended data Fig. 2G). GID4 interactors were associated with ribonucleoprotein complex biogenesis, chromatin binding, DNA binding, and spliceosomal complex, as well as ubiquitination and mitosis (Fig 4C). GID4 interactors were significantly enriched for GO terms associated with ribosomal RNA binding and processing, chromatin binding and organization, and the nucleus and nuclear lumen cellular compartments (Fig 4D). The GID4 interactome shares most similarity with the previously identified interactomes^29^ of other proteins localized to the nucleolus, nucleus, nucleoplasm, and chromatin (Fig 4E). Taken together, GID4 interacts broadly with distinct classes of nuclear proteins involved in RNA processing, chromatin, splicing, and transcription.

**Figure 4.**
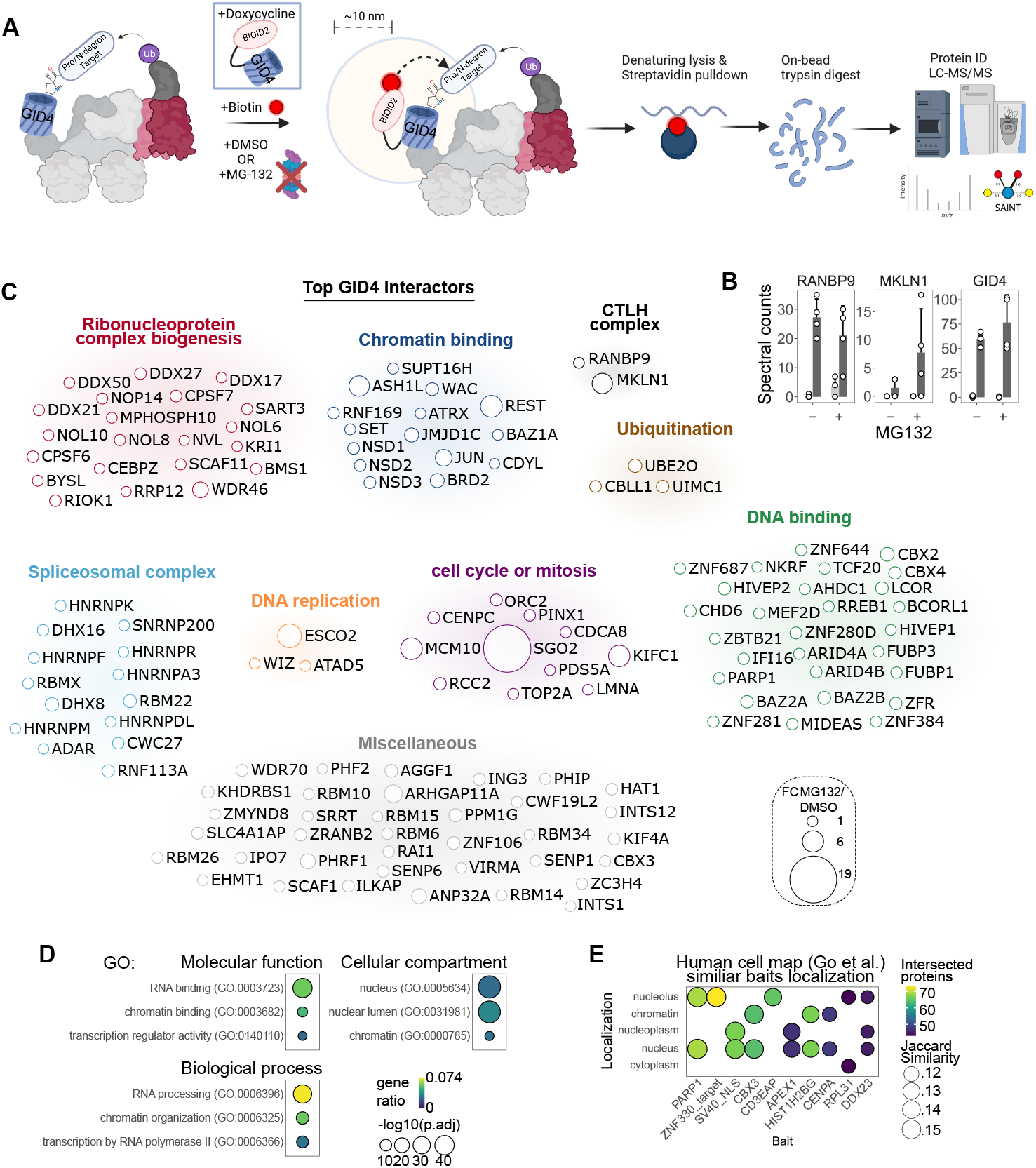
A) Schematic of GID4 proximity-dependent biotinylation workflow. Doxycycline-inducible BioID2-GID4 is expressed for 24h in the presence or absence of PFI-7. Upon induction, BioID2-GID4 binds to the endogenous CTLH complex and biotinylates proteins within approximately 10 nm radius. Biotinylated proteins are enriched before protein identification using liquid chromatography tandem mass spectrometry (LC-MS/MS). B) Spectral counts of CTLH complex members detected in BioID2-GID4. C) Top 133 GID4 interactors (saint probability [SP] > 0.9, spectral counts > 5) classified by functional annotation. The size of each circle represents fold change between MG132 and DMSO-treated samples. Larger circles contained more spectral counts in MG132. D) GO terms functional annotation of GID4 interactors (SP > 0.9). Gene ratio represents the fraction of genes in each GO category that were identified. P-values were adjusted using the Benjamini-Hochberg method. E) Top 10 most similar baits to GID4 found in the human cell map database^29^. Jaccard similarity represents the percentage overlap between sets of interacting proteins.

### RNA helicases DDX21 and DDX50 interact with GID4 via Pro/N-degron pathway

Since human substrates of the Pro/N-degron pathway have remained elusive, we determined the GID4 interactors that specifically bind via the GID4 substrate binding pocket by performing GID4 proximity-dependent biotinylation in the presence of proteasome inhibitor and in the presence and absence of PFI-7. Biotinylated proteins were quantified by LC-MS/MS in three groups; i) cells expressing BioID2 treated with DMSO (vehicle), ii) cells expressing BioID2-GID4 treated with DMSO, iii) cells expressing BioID2-GID4 treated with PFI-7. Treatment with PFI-7 broadly reduced BioID2-GID4 interactions, with the number of high-confidence GID4 interactors decreasing from 77 to 13 (SP ≥ 0.9, Fig. 5A, Supp. Table 3). Samples clustered by treatment group in UMAP and principal component (PCA) analyses (Fig. 5B). Interestingly, PFI-7 treatment of BioID2-GID4 samples caused them to cluster closer to BioID2 alone control samples than to DMSO-treated BioID2-GID4 cells (Fig. 5B). Six GID4 interactors were maintained, and a single interactor was gained after PFI-7 treatment (Fig. 5C). We reasoned that GID4 interactors lost after PFI-7 treatment would represent candidate targets of the Pro/N-degron pathway. Overall, there were 45 high-confidence interactors in DMSO-treated BioID2-GID4 cells that were significantly depleted in PFI-7 treated samples and fell below the medium confidence threshold including DDX17, DDX21, DDX50, and KIN, among others (SP BioID2-GID4 DMSO ≥ 0.9, BioID2-GID4 PFI-7 < 0.6, Fig. 5C). Functional annotation of GID4 interactors lost upon PFI-7 treatment revealed an enrichment of GO terms associated with nucleic acid binding, ribosome biogenesis, and the nucleus (Fig. 5D).

**Figure 5.**
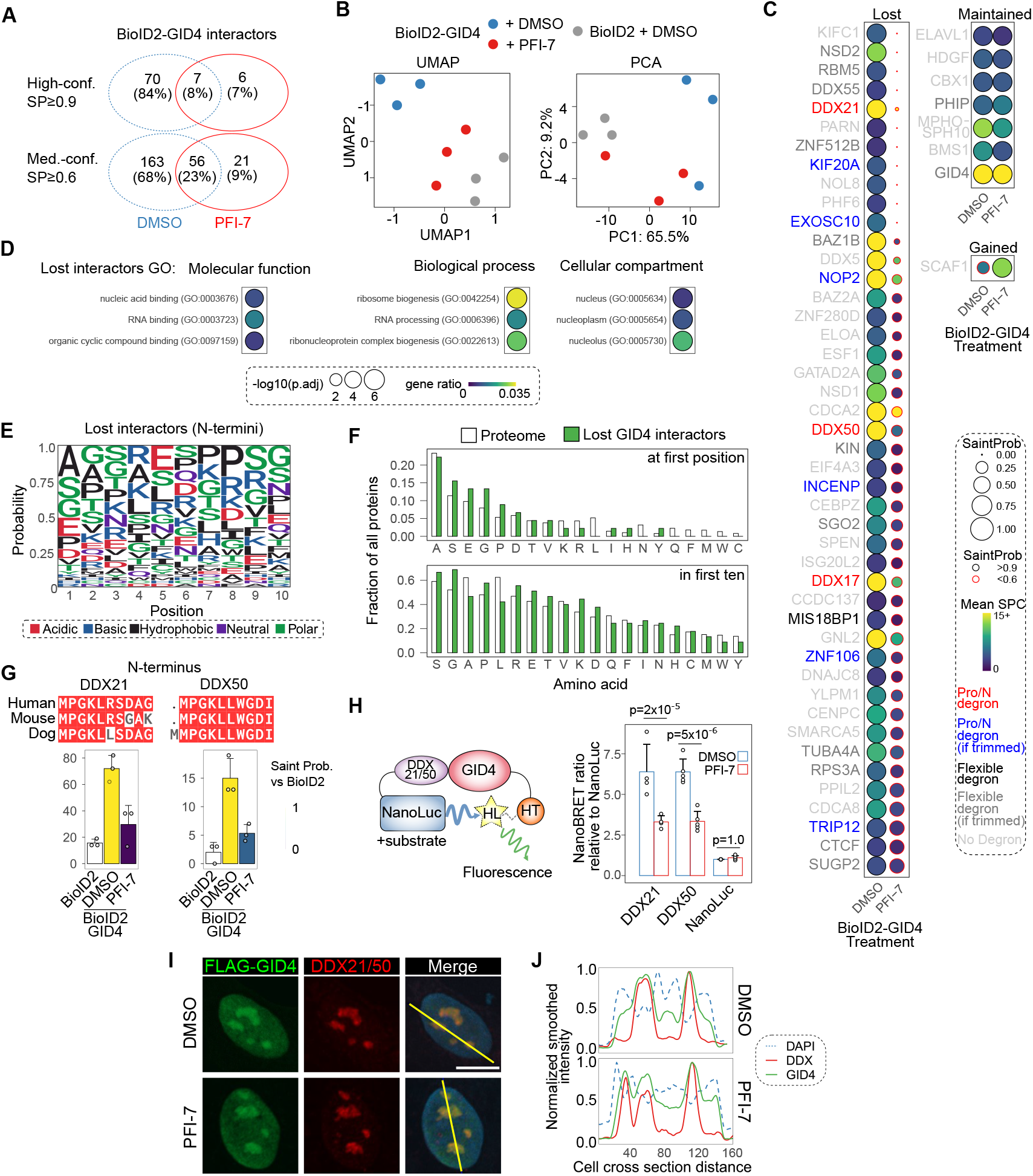
A) Numbers of high and medium confidence BioID2-GID4 interacting proteins identified in DMSO and PFI-7 treated samples. B) Clustering of BioID2 and BioID2-GID4 samples by uniform manifold approximation and projection (UMAP) and principal component analysis (PCA). Clustering was done on GID4-BioID log2-normalized area intensities. Missing or zero data for each protein was replaced with half of the lowest detected value for that protein. PCA was done on all proteins with saint probability (SP) > 0.6 in BioID2-GID4 condition (217 proteins). UMAP was done on proteins with SP > 0.6 in BioID2-GID4 and a difference in SP > 0.9 between BioID2-GID4 and BioID2-GID4 + PFI-7 (11 proteins). C) GID4 interactors that were lost (DMSO SP>0.9, PFI-7 SP<0.6), maintained (DMSO SP>0.9, PFI-7 SP>0.9), and gained (DMSO SP<0.6, PFI-7 SP>0.9) are shown. Circle size represents saint probability (SP), black outer circles color presents high confidence interactors (SP >0.9), red outer circles were not significant interactors (SP < 0.6). Inner circle color represents spectral counts intensity. Proteins are ranked by difference in SP between DMSO and PFI-7 treated samples. Protein names are colored according to their N-terminus. Red protein names contain a Pro/N-degron^10^ (N-terminal Pro followed by either of Ser/His/Thr/Ala), black contain a previously described flexible Pro/N-degron^24^ containing an hydrophobic amino acid (P/I/L/F/V) followed by a small side-chain residue (S/T/G/V/A), dark and light grey contain Pro/N-degrons or flexible degrons, respectively, within the first ten amino acids. D) GO terms functional annotation of GID4 interactors that were lost (DMSO SP>0.9, PFI-7 SP<0.6). Gene ratio represents the fraction of genes in each GO category that were identified. P-values were adjusted using the Benjamini-Hochberg method. E) Consensus sequence of ten N-terminal amino acids of GID4 interactors that were lost after PFI-7 treatment. F) Enrichment of different N-terminal amino acids at the first position or within the first ten amino acids of GID4 interactors that were lost after PFI-7 treatment (green bars) and the background in the proteome (white bars) *χ*2 tests df=19, p=0.810 (enrichments at first position), p=0.378 (enrichments in the first ten). G) N-terminal sequence conservation of DDX21 and DDX50 is shown. Spectral counts and Saint probability are shown for GID4-BioID2 vs BioID2 for DMSO and PFI-7-treated samples. H) Schematic of DDX21/50 GID4 PPI assay. Full-length C-terminal DDX21/50 NanoLuc fusion protein is shown interacting with N-terminal GID4-HaloTag protein to produce a fluorescence signal. Mean BRET Units normalized to NanoLuc alone in each experiment are shown for interactions between GID4 and DDX21-NL, DDX50-NL, NanoLuc. Red outlined bars were performed in the presence of PFI-7, blue outline bars were controls treated with vehicle (DMSO). One-way ANOVA revealed a significant interaction effect for protein and treatment on normalized BRET units (F(2)=19.46, p=6.83×10^−6^). Tukey’s honest significant difference test for multiple comparisons was done to identify differences between DMSO and PFI-7 treated samples (DDX21 adjusted p-value = 2.37×10^−5^, DDX50 adjusted p-value = 4.81×10^−6^, NanoLuc adjusted p-value = 1.00). I) Confocal imaging analysis of U2OS cells transduced with a doxycycline inducible lentiviral expression vector coding for N terminally FLAG-tagged GID4. FLAG-GID4 expression was induced for 24 h and is shown in green, DDX21/50 (antibody recognizes both endogenous proteins) is shown in red, and DAPI is shown in blue. Scale bar represents 15 μm. Representative images are shown from four independent experiments. J) Quantification of fluorescence signal intensity over a cross section of cell nuclei. A rolling average with a window size of 8 was used to smooth data and maximum intensities were scaled to 1.

To gain insights into the sequence specificity of the GID4-mediated interactions and to identify potential targets of the Pro/N-degron pathway in humans we examined the N-terminal sequences of PFI-7-dependent GID4 interactors. The N-termini of GID4 interactors DDX17, DDX21, and DDX50 matched a previously identified human GID4 Pro/N-degron sequence^10^ (N-terminal Pro followed by either of Ser/His/Thr/Ala^10^ ‘Pro/N degron’, Fig. 5C). One lost interactor matched a previously described more flexible Pro/N-degron^24^ containing a hydrophobic amino acid (P/I/L/F/V) followed by a small side-chain residue (S/T/G/V/A, ‘Flexible degron’, Fig. 5C). Other interactors harbored Pro/N-degrons or flexible degrons within the first ten amino acids (‘Pro/N-degron (if trimmed)’ and ‘flexible degron (if trimmed)’, Fig. 5C). The consensus sequence of the N-termini of high confidence lost interactors was distinct from both the canonical Pro/N-degron^10^, flexible degron^24^, and other described GID4 recognition sequence^25^ (Fig. 5E). Pro and Gly were slightly overrepresented in the N-terminal sequences of high-confidence lost interactors at position one and within the first ten amino acids, but this was not significantly different compared to the distribution of N-terminal amino acids in the proteome (Chi-squared test, df=19, p=0.810, 0.378 respectively, Fig. 5F).

GID4 interactors DDX21 and DDX50 that were lost upon PFI-7 treatment bear N-termini that are evolutionarily conserved and match the canonical Pro/N-degron. DDX21 was the most highly enriched high-confidence GID4 interactor by spectral counts (BioID2-GID4 DMSO mean spectral counts = 72, Supplementary Table 3), and both proteins interacted significantly less with GID4 after PFI-7 treatment (SP < 0.31, Fig. 5G). The interaction between GID4 and DDX21 and DDX50 was confirmed by an orthogonal NanoBRET PPI assay in live cells utilizing full-length C-terminally NanoLuc-tagged DDX21 or DDX50 and HaloTag-GID4 (Fig. 5H). Importantly, this interaction was blocked by PFI-7 treatment (Fig. 5H). FLAG-GID4 colocalized with DDX21/DDX50 in the nucleolus, and this colocalization was preserved after PFI-7 treatment (Fig. 5I, J), indicating that specific GID4-DDX21/50 interactions were blocked despite their nuclear colocalization being maintained. Furthermore, RANBP9 interacted with DDX21/50 in co-immunoprecipitation experiments and the interaction was reduced in ARMC8 knock-out cells (Extended data Fig. 3). Overall, GID4 specifically binds to proteins associated with ribosomal biogenesis including Pro/N-degron-bearing nucleolar RNA helicases DDX17, DDX21, and DDX50, and PFI-7 blocks these interactions.

### GID4 regulates protein levels of RNA helicases

In *Saccharomyces cerevisiae* GID4 is responsible for polyubiquitylating and degrading gluconeogenic enzymes Fbp1, Icl1, Mdh2^9^. However, the role GID4 plays in proteome regulation in humans is not clear. Since GID4 interacted with dozens of proteins in proximity-dependent biotinylation experiments, next we undertook a proteomics survey to determine the impact of GID4 overexpression on the proteome in the presence and absence of PFI-7 (Fig. 6A). Overall, 6068 proteins were quantified across all samples (per sample mean = 5427, standard deviation = 63, minimum quantified proteins = 5331, Fig. 6B). Protein levels of CTLH complex members were stable across treatments with the exception of GID4, which was significantly higher in BioID2-GID4 cells compared to BioID2 cells (Fig. 6C). Comparing across all treatment conditions, 427 proteins exhibited differential abundance (FC > 1.5, adjusted p-value < 0.05). Unsupervised hierarchical clustering of proteomics samples based on differentially abundant proteins revealed strong segregation based on proteasome inhibition and GID4 overexpression, with less robust clustering observed between PFI-7 and DMSO-treated samples (Fig. 6D). Principal component analysis revealed that proteasomal inhibition was the dominant driver of differential protein levels between groups (PC1, 50.4% of variance for all samples, 61.2% for BioID2-GID4 alone, Fig. 6E), with GID4 overexpression representing the second largest effect (PC2, 17.3% of variance for all samples). PFI-7 treatment led to distinct clustering of BioID2-GID4 samples that was reduced when proteasome function was inhibited (PC2, 9.9% of variance for BioID2-GID4, Fig. 6E). Differentially abundant proteins clustered broadly into four groups (Fig. 6D). Proteins that increased upon proteasomal inhibition were located in Cluster 3. GID4 overexpression decreased average levels of proteins within Cluster 1, while increasing levels of proteins in Cluster 4 (Fig. 6D, F). Cluster 2 contained proteins that decreased on average after PFI-7 treatment of BioID-GID4 cells, and this change was blocked by proteasomal inhibition (Fig. 6D, F). PFI-7 treatment led to a greater number of significantly changed proteins in cells with proteasomal function intact (60 differential proteins, FC > 1.5, adjusted p-value < 0.05) compared to cells treated with proteasome inhibitor (15 differential proteins) (Fig. 6G). PFI-7 treatment significantly increased levels of DHX40 and DICER1, while decreasing others including KIN and CHAF1A (Fig. 6G, H). GID4 overexpression altered levels of DDX39A and nucleolar protein IFI16 that were both increased, and EIF4A2 and LMNB2 that both decreased (Fig. 6G, H). There were similar numbers of protein level changes after GID4 overexpression with proteasomal function intact (55 proteins FC > 1.5, adjusted p-value < 0.05) or inhibited (62 proteins FC > 1.5, adjusted p-value < 0.05) (Fig. 6G). DDX21 and DDX50 protein levels were consistent irrespective of GID4 overexpression or PFI-7 treatment (Fig. 6G, H). PFI-7-dependent proteins included four that were identified as medium confidence GID4 interactors (BioID2-GID4 SP ≥ 0.6) including KIN and CHAF1A, and two that contained Pro/N-degrons, HMGCS1 and ASAH1 (Fig. 6H, Extended data Fig. 4A). Eight proteins altered after GID4 overexpression were identified as GID4 interactors including DDX39A and IFI16, and three contained Pro/N-degrons (Fig. 6H, Extended data Fig. 4B). Taken together, despite disrupting their interactions with GID4, protein levels of DDX21 and DDX50 remain similar after GID4 inhibition with PFI-7 while significant changes are observed in levels of other proteins including DHX40 and DICER1.

**Figure 6.**
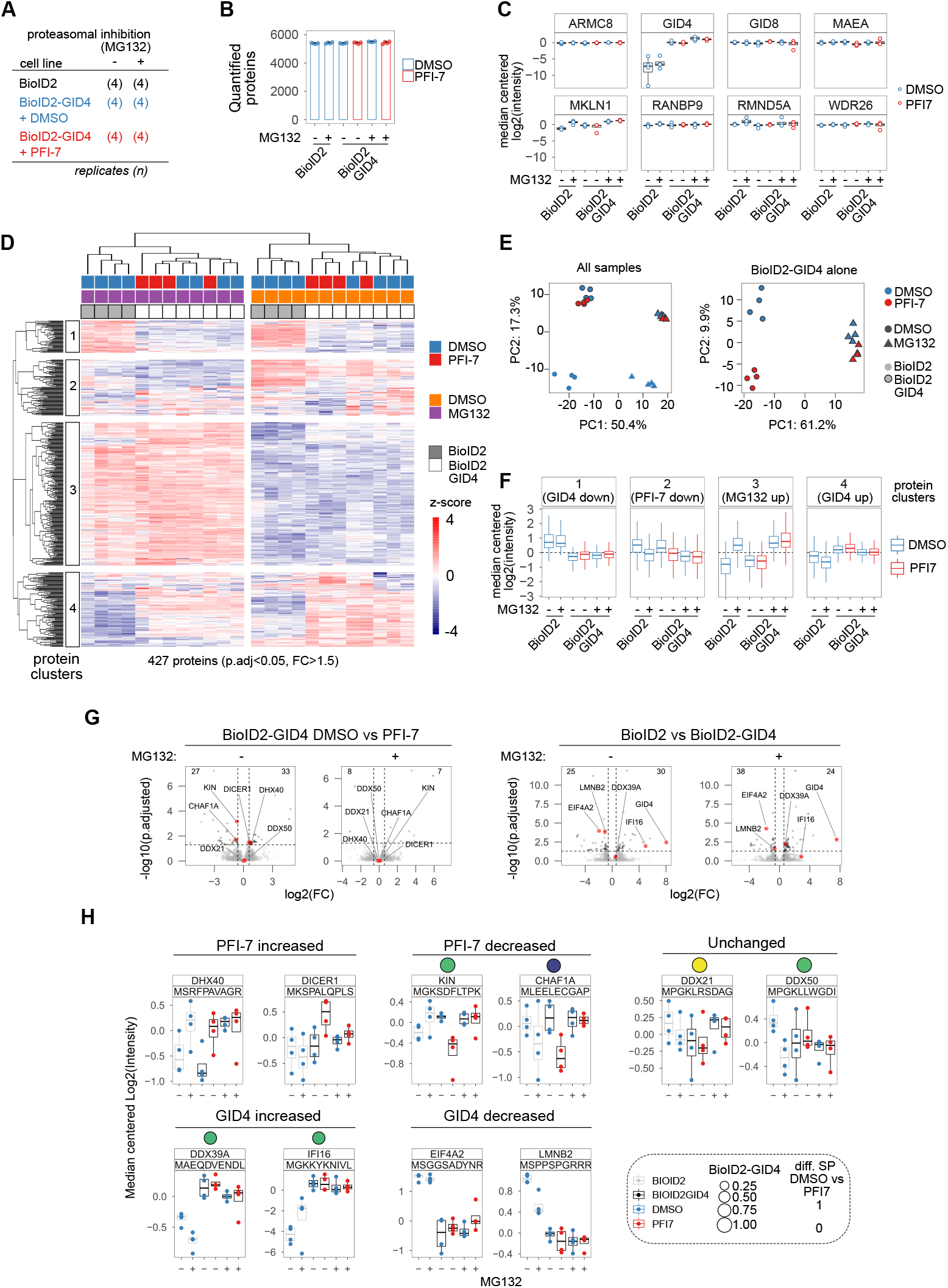
A) Proteomics samples with number of replicates and treatment conditions shown. B) Number of quantified proteins detected in each sample. C) Protein abundances of CTLH complex members. Median-centered log2 intensities are shown, with blue dots indicating DMSO-treated samples and PFI-7-treated samples shown in red. D) Heatmap of 427 differentially abundant proteins (adjusted p-value < 0.05, fold change > 1.5, statistics derived from DEP package, see methods). Each row represents one protein and samples are represented in columns Protein abundance is shown as z-scores quantified across rows. Hierarchical clustering of samples and proteins is shown. E) Principal component analysis of all samples (left) and BioID2-GID4 samples aline (right). DMSO-treated samples are indicated in blue and PFI-7-treated samples are shown in red. Samples treated with proteasome inhibitor (MG132) are indicated by triangles, and control samples are shown as circles. Shapes with a black outline represent BioID2-GID4 samples, while BioID2 alone samples have no outer shape color. F) Median protein abundance for proteins in each cluster for each treatment group. DMSO-treated samples are shown in blue and PFI-7 treated samples are shown in red. G) Volcano plot of all proteins with log2 fold change shown on the x-axis and −log10 adjusted p values shown on the y axis. The number of significantly changed proteins is indicated in the top corner. Specific proteins of interest are labeled and indicated by red dots. H) Median centered log2 intensities for specific proteins that were changed by PFI-7, GID4, or unchanged by either. First ten amino acids of N-termini are indicated below protein names. Grey boxplot outlines represent BioID2 alone samples, black boxplot outlines represent BioID2-GID4 samples. Blue dots represent DMSO-treated samples and red dots indicate PFI-7 treated samples. Circles plotted above each protein name represents enrichment in the BioID2-GID4 experiment indicated in Figure 5. Proteins with no circles above were not quantified in BioID2-GID4.

## Discussion

PFI-7 is a chemical probe targeting GID4, the substrate receptor of the CTLH ubiquitin ligase complex. As a high-quality chemical probe^30–33^ PFI-7 is potent (Kd of 79 ± 7 nM by SPR) and active in cells at 1 μM (NanoBRET GID4 MPGLWKS-NL PPI IC50 = 0.57 ± 0.05 μM). PFI-7 showed no evident cytotoxicity and no off-target activity against a panel of proteins such as kinases, GPCRs and other drug safety targets. The cellular activity and specific target engagement by PFI-7 is evident from the robust changes observed in GID4 interactions and proteome level regulation. Since the E3 ligase activity of the human CTLH complex has only recently been demonstrated^18,20^, and it is unclear what role GID4-mediated substrate recruitment and proteasomal degradation plays in the array of fundamental cellular processes the complex has been implicated in^12^ we analyzed global proteomics and the interactome of GID4 in the presence and absence of PFI-7 to probe the function of GID4 and the CTLH complex. We demonstrate that several proteins such as RNA helicases DDX21 and DDX50 are recognized by GID4 but are not substantively regulated at the protein level. This suggests that CTLH-GID4 may have degradative and non-degradative functions highlighting potential challenges for employing the CTLH complex for targeted protein degradation^8,26^. Thus, the unique advantages afforded through use of a potent and selective chemical probe provides new insights into GID4 and CTLH complex.

Our proximity-dependent biotinylation analysis defined the first GID4 interactome, complementing the rapidly growing numbers of interactomes for human proteins^29,34,35^. We demonstrate a strong tendency for GID4 to interact with nuclear and nucleolar proteins, which is supported by the immunofluorescence subcellular localization of exogenously expressed GID4. As the CTLH complex has been reported to form nuclear-specific complexes^36^, GID4 may be recruiting the CTLH complex to targets in the nucleus and nucleolus. One caveat of traditional proximity-dependent biotinylation studies is that both direct and indirect interactors within approximately 10 nm radius will be biotinylated^37,38^. PFI-7, however, facilitated the identification of proteins, or their interactors, that bind directly via the GID4 substrate binding pocket. Interactors that were lost upon PFI-7 treatment included RNA helicases DDX17, DDX21, and DDX50 that have previously been reported to associate with the CTLH complex^16,18,34,36^, and that we now show are GID4-dependent interactors. Indeed, many of the direct GID4 interactors are nucleolar proteins including DDX21^39^, DDX50^40^, and KIN^41^, among others. This points to possible nucleolar-specific functions of the CTLH complex that have been suggested previously^36^ and remain to be elucidated further.

Several of the nucleolar RNA helicases that interact with GID4 bear consensus Pro/N-degron sequences that agree with previously identified GID4 recognition sequences^10^ including DDX17, DDX21, and DDX50. Interestingly, many other interactors did not contain the canonical Pro/N-degron. There are several plausible explanations for this. Firstly, it has been suggested that the sequences GID4 binds to may be flexible and tolerate non-Pro hydrophobic residues^24,25^. In addition, specific N-terminal aminopeptidases were recently shown to trim protein N-termini to allow recognition of internal Pro/N-degrons in S cerevisiae^42^. The aminopeptidases identified are conserved in humans^42^ suggesting that some of the GID4 interactors identified that lack N-terminal recognition sequences might have been processed by aminopeptidases to reveal internal sequences that are recognized by GID4. Also, interactors might engage GID4 indirectly via intermediary proteins that carry conventional GID4 recognition sequences. The GID4 interactors we describe will inform future work to elucidate the seemingly diverse GID4-mediated recruitment mechanisms of the CTLH complex.

In addition to providing new insights into GID4 interactors, using PFI-7 we also characterized proteome level regulation by GID4 for the first time. We show that GID4 plays a role in regulating the cellular abundance of dozens of proteins including RNA helicase DHX40 and ribonuclease DICER1. Since GID4 antagonism with PFI-7 increased the levels of these proteins and this effect was blocked by proteasomal inhibition, this argues that GID4 is required for proteasomal degradation of DHX40 and DICER1. However, neither protein appears to interact with GID4 in our proximity-dependent biotinylation experiments, either suggesting that they do not directly interact with GID4, that their interaction topology is inconsistent with biotinylation, or that substrate lysines are not available to be biotinylated. Since we found that GID4 binds to several RNA helicases, a protein family known to facilitate multivalent interactions^43^, it is plausible that DHX40 and DICER1 might be binding to GID4 in complexes containing multiple RNA helicases. In support of this, DICER1 is a ribonuclease of the helicase family that interacts with an array of RNA binding proteins including RNA helicase DDX17^44,45^. DDX17 is a GID4 interactor lost upon PFI-7 treatment, bears a canonical N-terminal GID4-recognition sequence (PTGFVAPIL), and has previously been shown to interact with CTLH RING-domain subunit MAEA^34^. Similarly, DHX40 is a member of the DEAH-box family of RNA helicases^46^ that has previously been identified as an interactor of DDX23^29^ and DDX24^47^, both of which we identified as GID4 interactors (SP DDX23 = 0.67, SP DDX24 = 1.0). Together, this suggests that protein level regulation of specific targets might occur via indirect interactions mediated by intermediary proteins such as RNA helicases.

Intriguingly, PFI-7 treatment decreased levels of several proteins including KIN. KIN is a DNA and RNA binding protein implicated in ribosome biogenesis whose interactome shares several interactors with GID4, including DDX50^41^. There are several plausible explanations for how GID4 antagonism by PFI-7 might lead to decreased protein abundance. One intriguing possibility is that PFI-7 could be acting as a molecular glue^48,49^, leading to neo-substrate recruitment to GID4, polyubiquitylation by the CTLH complex and proteasomal degradation. This hypothesis is unlikely since GID4 interactions were lost broadly upon PFI-7 treatment, with only one interactor showing significantly increased interactions with GID4 upon PFI-7 treatment (SCAF1). This argues that PFI-7 likely does not act as a molecular glue. Alternatively, PFI-7-induced decreases in protein abundance could imply that certain interactions with GID4 might facilitate protein stabilization rather than degradation. GID4-mediated stabilization could occur if the CTLH complex mediates interactions between a target and a deubiquitinating (DUB) enzyme which may help sculpt ubiquitin chain configurations such that they result in activities other than degradation. Candidate DUBs for this include USP7 (a GID4 interactor, SP = 0.64), USP11 which interacts with CTLH complex via RANBP9 to stabilize Mgl-1^50^, or USP12, USP42, and USP46, which have all been reported to interact with the CTLH complex^28,51,52^. A third option is that GID4 antagonism induces transcriptional changes leading to reduced protein levels. However, the observed decreases in protein levels after PFI-7 treatment were dependent on the proteasome, showing that GID4 and proteasomal function are both involved in regulating levels of these proteins and arguing against this being solely a transcriptional effect.

While the abundances of dozens of proteins were altered by PFI-7 treatment, the majority of GID4 interactors were not affected. Protein-level regulation of DDX21 and DDX50 was absent, for instance, suggesting that their interaction with GID4 might impart other functions rather than polyubiquitylation-mediated degradation. In addition to proteasomal degradation, ubiquitylation is also associated with altered protein interactions and localization, among other functions^53^. In agreement with our identification of non-degradative GID4 interactions, ubiquitylation of the glycolysis enzymes PKM2 and LDHA was recently shown to be dependent on the CTLH complex, but the proteins were not degraded^23^. The ubiquitin linkages conjugated by E3 ubiquitin ligases depend on their cognate E2 enzymes^53^. The only E2 enzyme implicated so far in substrate ubiquitylation by the CTLH complex is UBE2H^18^. We observed significant interactions between GID4 and the unique E2/E3 hybrid enzyme, UBE2O. UBE2O is associated with monoubiquitylation of an array of substrates including ribosomal proteins^54^ and chromatin associated proteins^55^, and has previously been shown to co-fractionate with CTLH complex member WDR26^28,56^. This suggests that GID4 interactors such as DDX21/50 may be monoubiquitylated by CTLH/UBE2O to modulate their activity instead of being polyubiquitylated as a degradation signal. Alternatively, it is possible that DDX21/50 interact with GID4 in isolation from the rest of the CTLH complex, or with CTLH subcomplexes that do not support ubiquitylation activity. However, we found that the interaction between DDX21 and RANBP9 was lost after disrupting GID4 anchoring to the CTLH complex by knocking-out ARMC8^14^. Moreover, other CTLH complex members (MKLN1, RANBP9) interacted with GID4 in our proximity-dependent biotinylation experiments, showing that DDX21/50 do not interact with GID4 in isolation. Together, these data and our new results using PFI-7 suggest likely regulatory rather than degradative interactions between CTLH and its substrates.

PFI-7 facilitated the identification of dozens of GID4 interacting proteins, several of which are not substantively regulated at the protein level while other proteins were dependent on GID4 for protein-level regulation, possibly mediated through indirect interactions with the CTLH complex. It is clear that there is much still to learn about the fundamental biology of the mammalian CTLH complex. Going forward, PFI-7 will be a valuable research tool to study the CTLH complex and its substrate receptor GID4. Our findings inform future efforts to employ GID4 and the CTLH complex as an E3 ligase in targeted protein degradation.

## Methods

### Expression, purification of biotinylated GID4 and structure determination

The GID4 protein was purified as reported previously^10^. X-ray diffraction data for GID4 + PFI-7 was collected at 100K at beamline 24ID-C of Advanced Photon Source (APS), Argonne National Laboratory. The data were processed using XDS (Kabsch, W. (2010a). XDS. Acta Cryst. D66, 125-132.) and the HKL-3000 suite (Otwinowski Z, Minor W. Meth Enzymol 1997;276:307–326), respectively, and the structure was solved by molecular methods using PDB 6WZZ as a search template with the program PHASER (McCoy, A.J., Grosse-Kunstleve, R.W., Adams, P.D., Winn, M.D., Storoni, L.C., & Read, R.J., “Phaser crystallographic software”, J. Appl. Cryst. 40, 658-674 (2007)). REFMAC (Murshudov GN, Vagin AA, Dodson EJ. Acta Crystallogr D Biol Crystallogr 1997;53:240–255) and BUSTER (Bricogne G, Blanc E, Brandl M, Flensburg C, Keller P, Paciorek P, Roversi P, Sharff A, Smart O, Vonrhein C, Womack T (2010). BUSTER version 2.9. Cambridge, United Kingdom: Global Phasing Ltd.) were used for structure refinement. Graphics program COOT (Emsley P, Cowtan K. Acta Crystallogr D Biol Crystallogr 2004;60:2126–2132) was used for model building and visualization.

### Analysis of binding by Surface plasmon resonance (SPR)

In vitro binding analyses by SPR were carried out using a Biacore T200 (GE Health Sciences Inc.) instrument at 25 °C. HBS-EP+ buffer (10 mM HEPES pH 7.4, 150 mM NaCl, 3 mM EDTA, 0.05% Tween-20) was used for all the experiments. The N-terminally in vivo Biotinylated-GID4 (aa 116-300) was immobilised to approximately 2100 response units (RU) on a flow cell of a Streptavidin coated (SA) sensor chip (GE healthcare) according to manufacturer’s directions while another flow cell was left blank for reference subtraction and monitoring any non-specific bindings. The compounds were diluted in HBS-EP+ to yield a 1% final DMSO solution and were serially diluted (3-fold dilutions) in HBS-EP+ buffer containing 1% DMSO. The compounds were then flowed over the sensor chip using a MultiCycle Kinetics mode at 40 μL/min. Contact time (i.e. association phase) was 60 S and disassociation time was 120 S, which was followed by an additional 120 S of stabilization period under flow. HBS-EP+ buffer only (plus 1% DMSO) was used for blank injections; and HBS-EP+ buffers containing 0.5 to 1.5% DMSO were used for solvent correction injections. Binding constants were acquired from the double referenced (i.e. reference subtraction and blank injection subtraction) multicycle data using Biacore T200 Evaluation Software 3.1.

### Fluorescence polarization (FP)-based peptide displacement assay

The FP-based peptide displacement experiments were performed in 384-well polypropylene small volume black microplates (Cat. # 784209, Greiner) in a total volume of 15 μL per well at room temperature (i.e. 23 °C). Compounds were added to the reaction mixture (containing 5 μM GID4 (aa 116-300) and 40 nM of C-terminally FITC-labelled PGLWKS peptide in buffer (50 mM Tris, pH 7.5, and 0.01% Triton X-100)), and were incubated for 30 min at room temperature. Final DMSO concentration was 1.5%. The resulting FP signals were measured using a BioTek Synergy 4 (BioTek, Winooski, VT) at the excitation and emission wavelengths of 485 nm and 528 nm, respectively. The obtained FP values were blank subtracted and are shown as the percentage of control. All the experiments were carried out in triplicate (n=3) and the presented values are the average of replicates ± standard deviation. FP data were visualized using GraphPad Prism software 8.0 (La Jolla, CA).

### Cell culture

Cell lines were cultured according to standard aseptic mammalian tissue culture protocols in 5% CO2 at 37 °C HEK293 and HeLa cells were cultured in DMEM supplemented with 10% fetal bovine serum (FBS) and 100 U/ml penicillin and 100 μg/ml streptomycin. HeLa cells transiently transfected with the BioID2 plasmid constructs or HA-GID4 was done using jetPRIME transfection reagent (Polyplus-transfection, Illkirch-Graffenstaden, France), following the manufacturer’s instructions, and either harvested after 24 h for immunoblotting or fixed and permeabilized for immunofluorescence. For generation of T-REx™-293 inducible BioID2 cell lines, 1.5ug of DNA (1350 ng of pOG44 plasmid and 150ng of pCDNA5/FRT/TO/HA-myc-BIOID2-GID4 or pCDNA5/FRT/TO/HA-myc-BIOID2) was used to transfect 1 well of 6 well plate of Flp-In™ T-Rex™ HeLa doxycycline-inducible cells (generated by Dr. Arshad Desai’s laboratory, San Diego, CA, USA). On day 3, cells were selected with 200 ug/ml Hygromycin and 3 ug/ml Blasticidin until Day 17 when colonies were collected. Expression was induced with 1 ug/ml Doxycycline. Nuclear and cytoplasmic fractionation of lysates was conducted exactly as previously described (Onea et al 2022).

### Plasmids

To generate BioID2-GID4, GID4 cDNA was digested from the pEZ-M06 GID4 plasmid^20^ using PmeI and EcoRI restriction enzymes and ligated into Hpa1 and EcoR1-digested MYC-BioID2-MCS (Addgene plasmid #74223). An HA tag was subcloned 5’ to the MYC tag in the BioID2-GID4 plasmid as well as for negative control BioID2 alone vector. For generation of T-REx™-293 inducible BioID2 cell lines, The HA-myc-BioID2-GID4 construct was cloned into pCDNA5/FRT/TO vector. Alphafold predicted structure was generated using google collab (https://colab.research.google.com/github/sokrypton/ColabFold/blob/main/beta/AlphaFold2_advanced.ipynb^58,59^. For NanoBRET PPI assays, GID4 coding sequence was cloned in frame into pNLF1-N for GID4-NanoLuc N-terminal fusion, into pNLF1-C for GID4-NanoLuc C-terminal fusion, or into pHTN-HaloTag vector for GID4-HaloTag N-terminal fusion using the In-Fusion HD Cloning kit (Takara). Pro/N-degron coding sequence was cloned into pNLF1-C for MPGLWKS-NanoLuc C-terminal fusion. All backbone vectors were obtained from Promega prior to cloning and sequence verified to confirm desired cloned constructs.

### NanoLuciferase bioluminescence resonance energy transfer (NanoBRET)

#### GID4 Tracer NanoBRET

The assay was performed essentially as described^60^. Full-length GID4 N or C-terminal NanoLuc-fusion was transfected into HEK293T cells using FuGENE HD (Promega, E2312) according to manufacturer’s instructions and proteins were allowed to express for 20 hours. Serially diluted PFI-7 and GID4-Tracer at a concentration of 1 μM were pipetted into white 384-well plates (Greiner 781207) using an Echo acoustic dispenser (Labcyte). The corresponding protein-transfected cells were added and reseeded at a density of 2 × 10^5^ cells/mL after trypsinization and resuspending in Opti-MEM without phenol red (Life Technologies). The system was allowed to equilibrate for 2 hours at 37°C/5% CO2 prior to BRET measurements. To measure BRET, NanoBRET NanoGlo Substrate + Extracellular NanoLuc Inhibitor (Promega, N2540) was added as per the manufacturer’s protocol. Filtered luminescence was measured on a PHERAstar plate reader (BMG Labtech) equipped with a luminescence filter pair (450 nm BP filter (donor) and 610 nm LP filter (acceptor)). Competitive displacement data were graphed using GraphPad Prism 9 software using a normalized 3-parameter curve fit with the following equation: Y=100/(1+10^(X-LogIC50)).

#### NanoBRET PPI assay

For NanoBRET PPI assays, HEK293T cells plated in ninety 96-well white plastic plates (Greiner) were transfected with 10 ng NanoLuc protein (N-terminally tagged with MPGLWKS, full length DDX21, or full length DDX50) and 50 ng N-terminally tagged HaloTag-GID4 using X-tremeGENE HP DNA Transfection Reagent (Sigma), following the manufacturer’s instructions. The next day, the media was replaced with DMEM/F12 (no phenol red) supplemented with 4% FBS, penicillin (100 U/ml) and streptomycin (100 μg/ml) in the presence or absence of compounds and HaloTag NanoBRET 618 Ligand (Promega) for 4 h. Next, NanoBRET NanoGlo Substrate (Promega) solution was added to each well. Donor emission at 450 nm (filter, 450 nm/band-pass 80 nm) and acceptor emission at 618 nm (filter, 610 nm/long-pass) was measured within 10 min of substrate addition using a CLARIOstar microplate reader (Mandel). mBU values were calculated by subtracting the mean of 618/460 signal from cells without a NanoBRET 618 Ligand × 1,000 from the mean of 618/460 signal from cells with a NanoBRET 618 Ligand × 1,000.

#### GID4-Tracer synthesis

*3-(5,5-difluoro-7-(1H-pyrrol-2-yl)-5H-5λ4,6λ4-dipyrrolo[1,2-c:2’,1’-f][1,3,2]diazaborinin-3-yl)-N-(2-(2-(2-(2-((2-((1s,4s)-4-(2-((4-methoxybenzyl)amino)acetamido)cyclohexyl)-1H-benzo[d]imidazol-6-yl) oxy) ethoxy) ethoxy) ethoxy) ethyl) propanamide*

A mixture of N-((1s,4s)-4-(6-(2-(2-(2-(2-aminoethoxy)ethoxy)ethoxy)ethoxy)-1H-benzo[d]imidazol-2-yl)cyclohexyl)-2-((4-methoxybenzyl)amino)acetamide (9.9 mg, 17 μmol), 2,5-dioxopyrrolidin-1-yl 3-(5,5-difluoro-7-(1H-pyrrol-2-yl)-5H-5λ4,6λ4-dipyrrolo[1,2-c:2’,1’-f][1,3,2]diazaborinin-3-yl)propanoate (6.0 mg, 14 μmol) and N,N-diisopropylethylamine (4.9 μL, 28 μmol) in anh. DMF (0.5 mL) was stirred at ambient temperature for 1 h and afterwards purified by prep. HPLC (H2O/ACN with 0.1% TFA) to provide the title compound (10 mg, 79%).

1H NMR (500 MHz, DMSO-d6): δ 11.43 (s, 1H), 9.13 (s, 2H), 8.30 (d, J = 7.3 Hz, 1H), 8.01 (t, J = 5.6 Hz, 1H), 7.63 (d, J = 9.0 Hz, 1H), 7.43 (s, 1H), 7.40 – 7.35 (m, 3H), 7.33 (d, J = 4.5 Hz, 1H), 7.27 (td, J = 2.7, 1.4 Hz, 1H), 7.20 (d, J = 2.3 Hz, 1H), 7.17 (d, J = 4.5 Hz, 1H), 7.12 (dd, J = 9.2, 2.2 Hz, 1H), 7.01 (d, J = 4.0 Hz, 1H), 6.99 – 6.95 (m, 2H), 6.36 – 6.31 (m, 2H), 4.18 – 4.13 (m, 2H), 4.08 (s, 2H), 3.99 – 3.93 (m, 2H), 3.81 – 3.77 (m, 2H), 3.75 (s, 3H), 3.65 – 3.48 (m, 11H), 3.42 (t, J = 5.9 Hz, 2H), 3.31 – 3.25 (m, 1H), 3.22 (q, J = 5.8 Hz, 2H), 3.13 (t, J = 7.8 Hz, 2H), 2.04 – 1.97 (m, 4H), 1.77 – 1.69 (m, 2H), 1.66 – 1.60 (m, 2H). 13C NMR (126 MHz, DMSO): δ 171.05, 164.21, 159.81, 156.14, 155.87, 150.22, 136.93, 132.99, 132.42, 131.72, 131.66, 126.73, 126.09, 124.38, 123.09, 122.88, 119.33, 116.10, 114.07, 111.50, 69.95, 69.81, 69.75, 69.58, 69.09, 68.85, 68.02, 55.19, 49.28, 46.34, 44.48, 44.44, 33.99, 33.77, 28.42, 28.39, 28.36, 28.34, 28.32, 24.61, 24.00, 23.97. MS (ESI): m/z calc. for [C_47_H_57_BF_2_N_8_O_7_+Na^+^]^+^ = 917.43, found = 917.45

### Proximity-dependent biotinylation (BioID2)

For biotinylation experiments, three (BioID2 experiment #1 (BE1): DMSO versus MG132) or two (BioID2 experiment #2 (BE2): PFI-7 treatments) 15 cm plates of each inducible HeLa cell lines were grown to 85–95% confluency and induced with 1 ug/mL doxycycline. Four hours later, cells were supplemented with 50 uM filter-sterilized biotin. For BE1, cells were also treated with 2 uM MG132 or equivalent volume DMSO, or for BE2, 2 uM MG132 and 10 uM PFI-7 or equivalent volume DMSO. Twenty-four hours later, cells were washed thoroughly in PBS, harvested and pellets were flash frozen and stored at −80. Cell pellets were lysed in 1300 uL RIPA lysis buffer on ice for 30 minutes with vortexing every 5 minutes (0.1% SDS, 0.5% sodium deoxycholate, 1% NP-40, 50 mM Tris-HCL, 150 mM NaCl, supplemented with Benzonase^®^ Nuclease (Sigma-Aldrich, cat# E1014) and protease inhibitors 0.2 mM phenylmethane sulfonyl fluoride (PMSF), 1 mM dithiothreitol (DTT), 1 g/mL leupeptin, 10 g/mL aprotinin, 1 g/mL pepstatin (inhibitors obtained from BioShop, Burlington, ON, Canada). Cell extracts were then sonicated using a Fisher Scientific Sonic Dismembrator, model 100; 30 × 1 second pulses at power level 2. Lysates were centrifuged 17, 968 xg 4 °C, 30 min, and quantified using using 660 nm protein quantitation assay (Thermo, #22660, #22663). A volume corresponding to 1.5 mg protein extract was incubated overnight with 50 uL Dynabeads (MyOne Streptavidin C1; Invitrogen). Protein-bound beads were collected and washed once with Strep-biotin wash buffer (50 mM Tris-HCL pH 8, 1% SDS (w/v), 150 mM NaCl) at RT, rotating for 5 min. Beads were then washed twice with RIPA lysis buffer, followed by three washes in TAP lysis buffer (10% glycerol, 0.1% NP-40, 2 mM EDTA pH 8, 50 mM HEPES pH 7.9, 100 mM KCl). Finally, beads were washed three times using 50 mM NH4HCO3 (ammonium bicarbonate, ABC) pH 8.0 solution before resuspension in ABC solution. Next, protein-bound beads were digested with 0.2 ug mass spectrometry-grade Lysyl Endopeptidase^®^ (Lys-C) (125-05061, Wako Pure Chemical Ind., Osaka, Japan) at 37 C for 2 h before digestion overnight with 1 ug Trypsin/Lys-C mix (V5071, Promega, Madison, WI, USA) at 37 C. The released peptides were then transferred over to a new tube and a 4 h digest using 0.5 ug mass spectrometry-grade trypsin (V5111, Promega) at 37 C for 4 hours was conducted. Reactions were then acidified to a final concentration of 1% TFA and spun at max speed. The supernatant was then applied to Pierce™ C18 Spin Tips, Cat# 84850 for desalting following the manufacturers protocol. The eluted peptides were speed vacuumed to near dryness then resuspended in 20 uL 0.1% FA. A volume corresponding to 500 ng peptides, as determined by BCA assay was then injected onto a Waters M-Class nanoAcquity HPLC system (Waters) coupled to an ESI Orbitrap mass spectrometer (Q Exactive plus, ThermoFisher Scientific). Buffer A consisted of mass spectrometry grade water with 0.1% FA and buffer B consisted of acetonitrile with 0.1% FA (ThermoFisher Scientific). All samples were trapped for 5 minutes at a flow rate of 5 μL/min using 99% buffer A and 1% buffer B on a Symmetry BEH C18 Trapping Column (5 mm, 180 mm × 20 mm, Waters). Peptides were separated using a Peptide BEH C18 Column (130 Å, 1.7 mm, 75 mm × 250 mm) operating at a flow rate of 300 nL/min at 35°C (Waters). Samples were separated using a non-linear gradient consisting of 1%-7% buffer B over 1 minute, 7%-23% buffer B over 59 minutes, and 23%-35% buffer B over 20 minutes, before increasing to 98% buffer B and washing. Full MS spectra were acquired in positive mode at R = 70 000 in the 400-1500 m/z mass range, 250 ms injection time, 3 × 106 ACG target. Top 12 peptides were selected for higher-energy collisional dissociation at R = 17 500 (ACG target: 2 × 105; injection time: 64 ms; loop count: 12; isolation width: 1.2 m/z; isolation offset: 0.5 m/z; normalized collision energy: 25; intensity threshold: 3.1 × 104; charge exclusion: unassigned, 1, 7, 8, >8; dynamic exclusion enabled 30 seconds). For BE1, data was searched using MaxQuant v1.5.8.3 using the Human Uniprot database (reviewed only; updated May 2017 with 42,183 entries). Missed cleavages were set to 3, cysteine carbamidomethylation (CAM) was set as a fixed modification and oxidation (M), N-terminal acetylation (protein) and deamidation (NQ) were set as variable modifications (max. number of modifications per peptide = 5), and peptide length ≥7. All other parameters were left at default. For BE2, data was searched using PEAKS Studio version 8.5 (Bioinformatics Solutions Inc., Waterloo, ON, Canada) using the Human Uniprot database (reviewed only; updated November 2019). Missed cleavages was set to 3 and semi-specific cleavage was enabled. Parent mass error tolerance was set to 20 ppm, fragment mass error tolerance was set to 0.8 Da. Protein and peptide False Discovery Rate (FDR) was set to 1%. Cysteine carbamidomethylation was set as a fixed modification while oxidation (M) and N-terminal deamidation (NQ) were set as variable modifications (maximum number of modifications per peptide = 5). Proteins were identified using a minimum of 2 unique peptide(s) with the FDR set to 1%. Spectral counts data were formatted for SAINTexpress analysis, a computational algorithm integrated into the REPRINT website (https://reprint-apms.org/). Analyzed data were visualized in cytoscape (14597658). GO terms were identified using the package goseq (1.42.0) in R (4.0.3) with p values adjusted using the Benjamini-Hochberg method and a threshold of adjusted p < 0.05. Comparison between GID4 interactome and the human cell map database was done at https://humancellmap.org/ against the entire database^29^.

### Global quantitative proteomics using label-free LC-MS/MS

For proteome sample preparation, MG-treated samples were the same extracts used in BE2. For non-MG samples, a separate 10 cm cell dish grown in parallel with BE2 treatments and replicates was treated with an equivalent volume of DMSO and collected and lysed at the same time as the BE2 extracts. Twenty-five microgram of protein lysate was reduced in 10 mM DTT for 25 minutes, alkylated in 10 mM iodoacetamide for 25 minutes in the dark, followed by methanol precipitation. The protein pellet was resuspended in 50 mM ABC and subjected to a sequential digest first with 250 ng of LysC (125-05061, Wako Pure Chemical Ind., Ltd., Japan) for 4 hours, then 500 ng of Trypsin/LysC (V5071, Promega, Madison, WI, USA) for 16 hours, followed by 500 ng of Trypsin (V5111, Promega) for an additional 4 hours. Digestions were incubated at 37°C at 600 rpm on a Thermomixer C (cat# 2231000667, Eppendorf, Hamburg, Germany). After the last digestion, samples were acidified with 10% formic acid (FA) to pH 3-4, centrifuged at 14 000 g for 5 minutes, then peptides desalted using Pierce™ C18 Spin Tips (Cat# 84850). Samples were then dried in a Speed vacuum, resuspended in 0.1% formic acid, and quantified by BCA assay. Approximately 500 ng of peptide sample was injected onto a Waters M-Class nanoAcquity UHPLC system (Waters, Milford, MA) coupled to an ESI Orbitrap mass spectrometer (Q Exactive plus, ThermoFisher Scientific). Samples were trapped for 5 minutes at a flow rate of 5 μL/min using 99% buffer A and 1% buffer B on a Symmetry BEH C18 Trapping Column (5 mm, 180 mm × 20 mm, Waters). Peptides were separated using a Peptide BEH C18 Column (130 Å, 1.7 mm, 75 mm × 250 mm) operating at a flow rate of 300 nL/min at 35°C (Waters). Proteome samples were separated using a non-linear gradient consisting of 1%-7% buffer B over 1 minute, 7%-23% buffer B over 179 minutes, and 23%-35% buffer B over 60 minutes, before increasing to 98% buffer B and washing. Full MS spectra were acquired in positive mode at R = 70 000 in the 400-1500 m/z mass range, 250 ms injection time, 3 × 106 ACG target. Top 12 peptides were selected for higher-energy collisional dissociation at R = 17 500 (ACG target: 2 × 105; injection time: 64 ms; loop count: 12; isolation width: 1.2 m/z; isolation offset: 0.5 m/z; normalized collision energy: 25; intensity threshold: 3.1 × 104; charge exclusion: unassigned, 1, 7, 8, >8; dynamic exclusion enabled 30 seconds). All MS raw files were searched in MaxQuant version 1.5.8.3 using the Human Uniprot database (reviewed only; updated July 2020). Missed cleavages were set to 3, cysteine carbamidomethylation (CAM) was set as a fixed modification and oxidation (M), N-terminal acetylation (protein) and deamidation (NQ) were set as variable modifications (max. number of modifications per peptide = 5), and peptide length ≥6. Protein and peptide FDR was left to 0.01 (1%) and decoy database was set to revert. Match between runs was enabled and all other parameters left at default. Protein groups were loaded into R (4.0.3, R Core Team (2020), https://www.R-project.org/) and reverse, potential contaminant, or proteins quantified in less than two out of four replicates of at least one condition were removed. Uniprot IDs were matched to gene symbols using biomaRt^61^ (2.46.3) and Uniprot. Mixed imputation was performed using DEP^62^ (1.12.0) as described in the package documentation. Testing for protein differential abundance was done using DEP with an adjusted p-value threshold of < 0.05, fold change > 1.5. Heatmap and hierarchical clustering was done using pheatmap (Raivo Kolde (2019), https://CRAN.R-project.org/package=pheatmap, 1.0.12), UMAP clustering was done using umap (0.2.7.0, Tomasz Konopka (2020), https://CRAN.R-project.org/package=umap). Principal Components Analysis was done using the stats package function prcomp (R 4.0.3).

### Immunoprecipitation

RanBPM immunoprecipitation in whole-cell extracts of BioID2-GID4 HeLa transfected cells was conducted exactly as previously described for RanBPM affinity purification mass spectrometry using DMP-crosslinked RanBPM antibody to dynabeads with 0.5 mg of protein extract, 5 ug RanBPM antibody (F1, sc-271727, Santa Cruz Biotechnology, Santa Cruz, CA, USA), and 20 uL Dynabeads Protein G (10004D, Invitrogen, Life Technologies, Burlington, ON, Canada) (Maitland et al 2021). Immunoprecipitation of HA-BioID2-GID4 with 5 ug of HA antibody (HA-7, H9658, Sigma-Aldrich), 20 uL of Dynabeads Protein G, and 0.5 mg of protein extract was conducted exactly as previously described (Maitland et al 2019).

### Fluorescence microscopy

HeLa BioID2-GID4 or BioID2 alone cells were seeded onto coverslips, fixed using 4% paraformaldehyde (PFA) 10 minutes 4 C, then permeabilized with 0.5% Triton X-100 for 15 min at room temperature (RT). Cells were blocked in 5% FBS then incubated overnight at 4 C with primary mouse antibody against HA (H9658, Sigma-Aldrich, Oakville, ON, Canada, 1:1000) and then with secondary Alexa 488 antibody against mouse (Invitrogen, Life Technologies, Burlington, ON, Canada, 1:1000). The coverslips were mounted using ProLong Gold containing 4,6-diamidino-2-phenylindole (DAPI) (Invitrogen). Cell images were taken with an Olympus BX51 microscope at 40× magnification and Image-Pro Plus (v5.0) software (Media Cybernetics, Inc., Bethesda,MD, USA). For microscopy of FLAG tagged GID4, 2×10^5^ U2Os cells were seeded in 12-well plates with 1 μg of doxycycline with 10 μM PFI-7 or DMSO control for 24 hours. Cells were fixed with 2% paraformaldehyde in 1× phosphate buffered saline (PBS) for 10 minutes and permeabilized with 0.1% Triton X-100 in 1× PBS for 5 mins at room temperature. Samples were then blocked with 2% bovine serum albumin (BSA; Sigma) in PBS-T (1× PBS and 0.1% Tween 20) for 1 hour at room temperature and incubated overnight at 4°C with primary antibodies staining for FLAG (Sigma Cat# F1804, 1:500) and DDX50 (Proteintech Cat# 10358-I-AP, 1:400). Cells were washed three times with 1× PBS, followed by staining with fluorescent anti-mouse (Alexa Fluor^®^ 488 Conjugate; Cell Signalling Technologies; # 44085, 1:400) and anti-rabbit (Alexa Fluor^®^ 594 Conjugate; Cell Signalling Technologies; #8889, 1:400) for 1 hour at room temperature. Unbound secondary antibodies were removed by three 1× PBS washes and coverslips were mounted onto slides using with Fluoroshield with DAPI (Sigma Cat#F6057). Images were acquired with Quorum Spinning Disk Confocal Microscope equipped with 405, 491, 561, and 642 nm lasers (Zeiss) and processed with Volocity software (Perkin Elmer) and ImageJ.

### Western blotting

For immunoblotting of experiments relating to BioID2-GID4, extracts were resolved by SDS-PAGE (10%) before transfer onto a polyvinylidene difluoride (PVDF) membrane and blocking in 5% skim milk in TBST solution. Membranes were hybridized overnight with the following antibodies: mouse anti-HA (H3663, Sigma-Aldrich, 1:5000), RanBPM (5M, 71-001, Bioacademia, Japan, 1:2000), Vinculin (E1E9V, Cell Signaling Technology, Danvers, MA, USA, 1:10000), SAP62 (1:1000; A-3, sc-390444, Santa Cruz Biotechnology), Muskelin (C-12, sc-398956, Santa Cruz Biotechnology, 1:1000), ARMC8 (E-1, sc-365307; Santa Cruz Biotechnology, 1:500), WDR26 (ab85962, Abcam, Cambridge, UK, 1:2000), RMND5A (1:2000; custom-made antibody from Yenzyme Antibodies), and MAEA (AF7288, R&D Systems, Minneapolis, MN, USA, 1:250). Biotinylated proteins were detected similarly: following transfer, PVDF membranes were blocked overnight in 0.5% fish gelatin in TBST solution and incubated for 1 h with horse radish peroxidase (HRP)-conjugated streptavidin (PierceTM High Sensitivity Streptavidin-HRP, ThermoFisher Scientific, Waltham, MA, USA, 1:20,000). Blots were imaged using the ClarityWestern ECL substrate (Bio-Rad, Hercules, CA, USA) and the Molecular Imager^®^ ChemiDocTM XRS system (Bio-Rad) with Image Lab (v6.0.1.).

## Data and code availability

The mass spectrometry data have been deposited to the ProteomeXchange Consortium via the PRIDE partner repository with the dataset identifier PXD038487. R scripts for analysis of proteomics data are freely available at https://github.com/d0minicO/GID4_analysis.

## Acknowledgements

This work was supported by Mitacs Elevate Postdoctoral Fellowship to D.D.G.O, funding from the Canadian Institutes for Health Research (CIHR) (MOP-142414 and PJT-169101 to C.S-P; FDN154328 to C.H.A.), and NSERC grant RGPIN-2021-02728 to J.M. D.B-L. is supported by NSERC grant 2021-03435 and CRS grant 25418. M.P.S is funded by the Deutsche Forschungsgemeinschaft (DFG, German Research Foundation), CRC1430 (Project-ID 424228829). We thank M. Robers and K. Riching from Promega for advising on the NanoBRET and target engagement assays. This research used resources of the Advanced Photon Source, a U.S. Department of Energy (DOE) Office of Science user facility operated for the DOE Office of Science by Argonne National Laboratory under Contract No. DE-AC02-06CH11357. Mass spectrometry analyses were performed on equipment funded by a grant from the Canada Foundation for Innovation to G.A.L. The Structural Genomics Consortium is a registered charity (no: 1097737) that receives funds from Bayer AG, Boehringer Ingelheim, Bristol Myers Squibb, Genentech, Genome Canada through Ontario Genomics Institute [OGI-196], EU/EFPIA/OICR/McGill/KTH/Diamond Innovative Medicines Initiative 2 Joint Undertaking [EUbOPEN grant 875510], Janssen, Merck KGaA (aka EMD in Canada and US), Pfizer and Takeda.

---

**Extended Data Figure 1.**
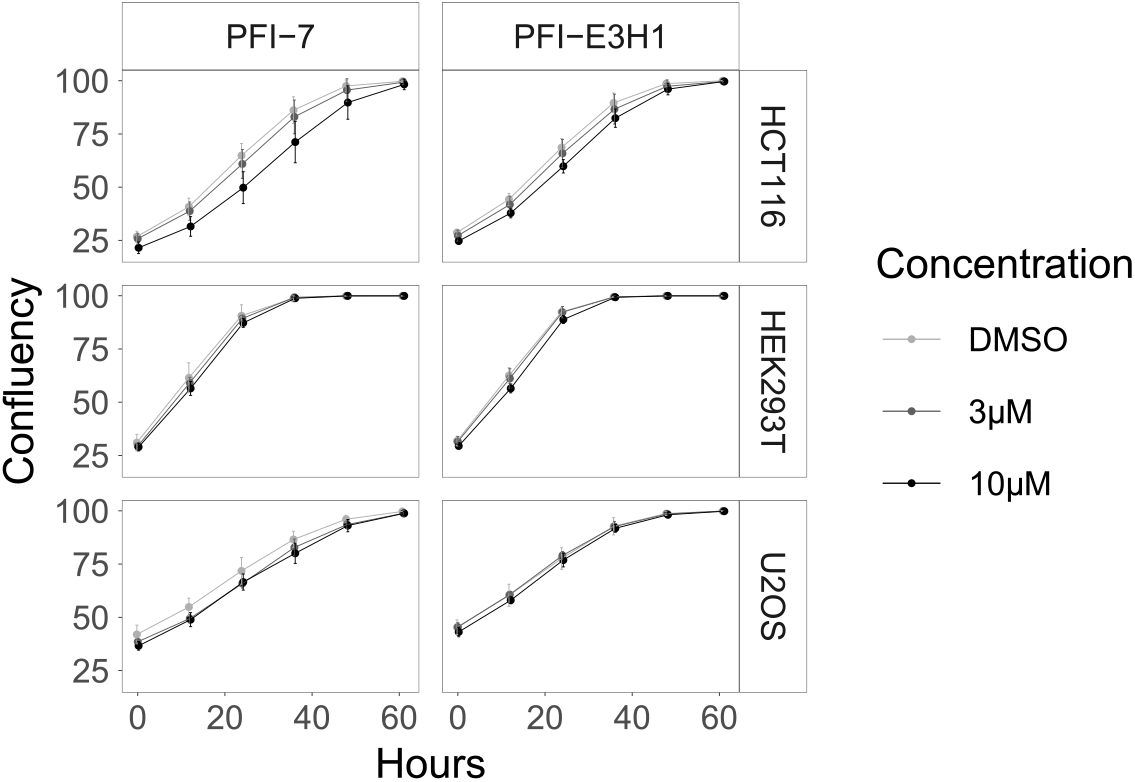
Cell growth curves for HCT116, HEK293T and U2OS cells treated with increasing concentrations of PFI-7 or PFI-E3H1 are shown over three days.

**Extended Data Figure 2.**
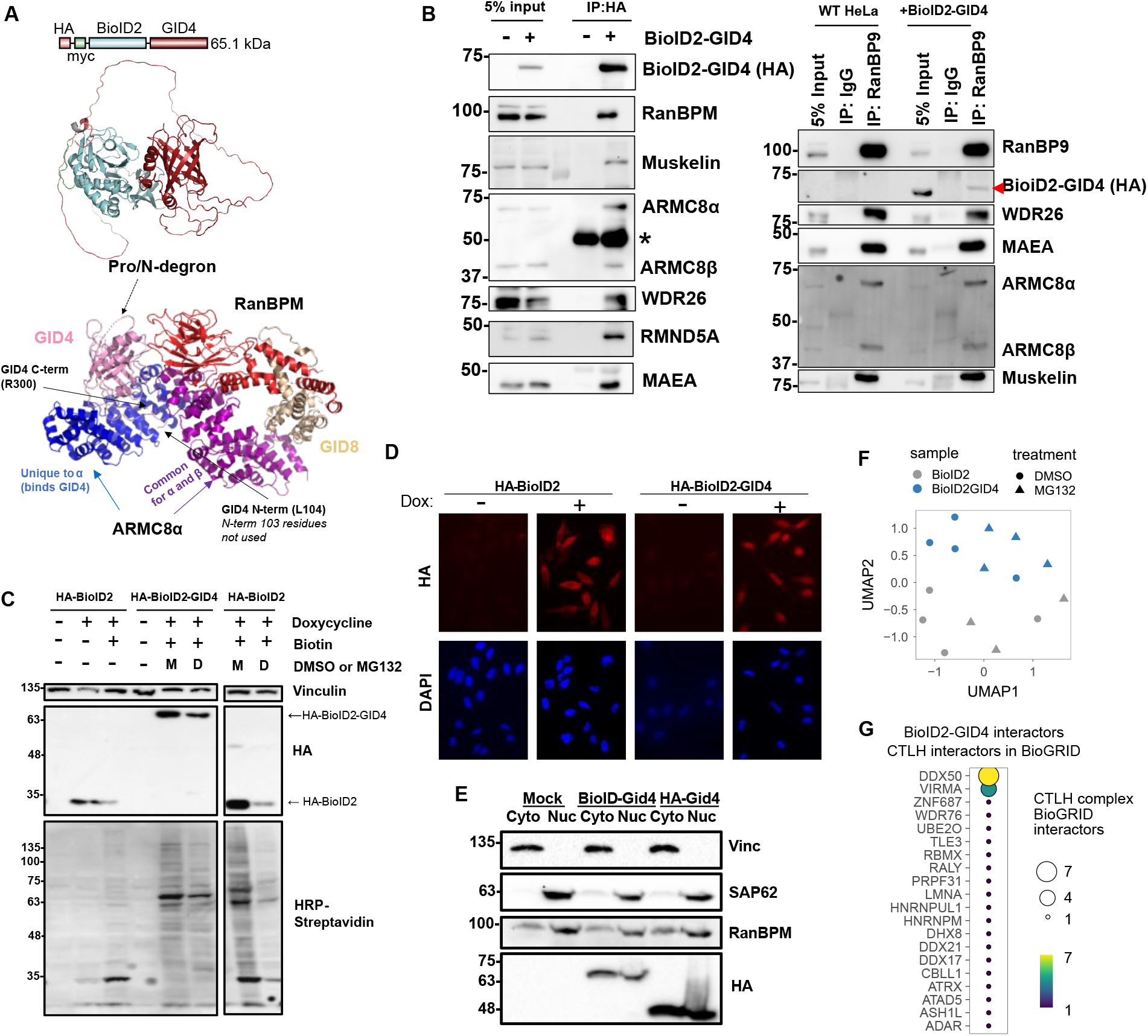
A) Upper, schematic and Alphafold2 predicted structure of GID4-BioID2 fusion protein. Lower, partial CTLH complex structure (PDB 7NSC) showing GID4 C-terminal anchor interacting with ARMC8α^57^. B) Immunoprecipitation experiments performed in HeLa cells. Left, HA-pull down of BioID2-GID4 fusion protein with immunoblotting to detect CTLH complex members. Right, immunoprecipitation of RanBP9 with immunoblotting to detect BioID2-GID4 (HA) and other CTLH complex members, *=heavy chain. C) Immunoblotting to detect BioID2-GID4 or BioID2 alone after doxycycline induction (top). Bottom, streptavidin-based detection of biotinylated proteins. D) Immunofluorescence imaging of HA in BioID2 and BioID2-GID4-expressing HeLa cells. HA signal is shown in red, DAPI is shown in blue. E) Nuclear and cytoplasmic fractionation of HeLa cells expressing BioID2-GID4 or HA-GID4 alone. Immunoblotting of Vinculin, SAP62, RanBP9, and HA is shown. F) Uniform Manifold Approximation and Projection (UMAP) analysis of proximity-dependent biotinylation samples. BioID2 and BioID2-GID4 cells are shown in gray and blue, respectively. MG132-treated samples are indicated as triangles, and samples treated with vehicle (DMSO) are shown as circles. UMAP was done on 196 high confidence GID4 interacting proteins (MG132 SP > 0.9) G) Overlap between high confidence GID4 interactors and interactors of the CTLH complex present in the BioGRID database^28^.

**Extended Data Figure 3.**
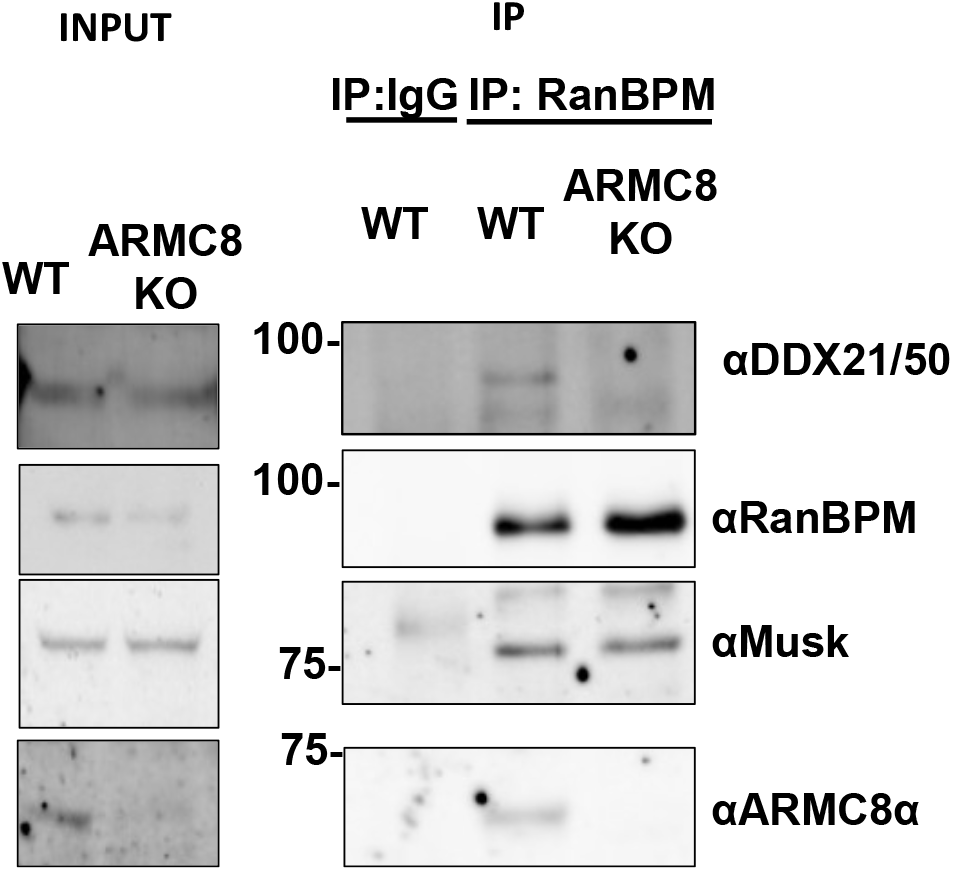
Immunoprecipitation of RanBP9 in wild type (WT) and ARMC8 knock-out (ARMC8 KO) HeLa cells. Immunoblotting was done for DDX21/50, RanBP9, Muskelin and ARMC8α.

**Extended Data Figure 4.**
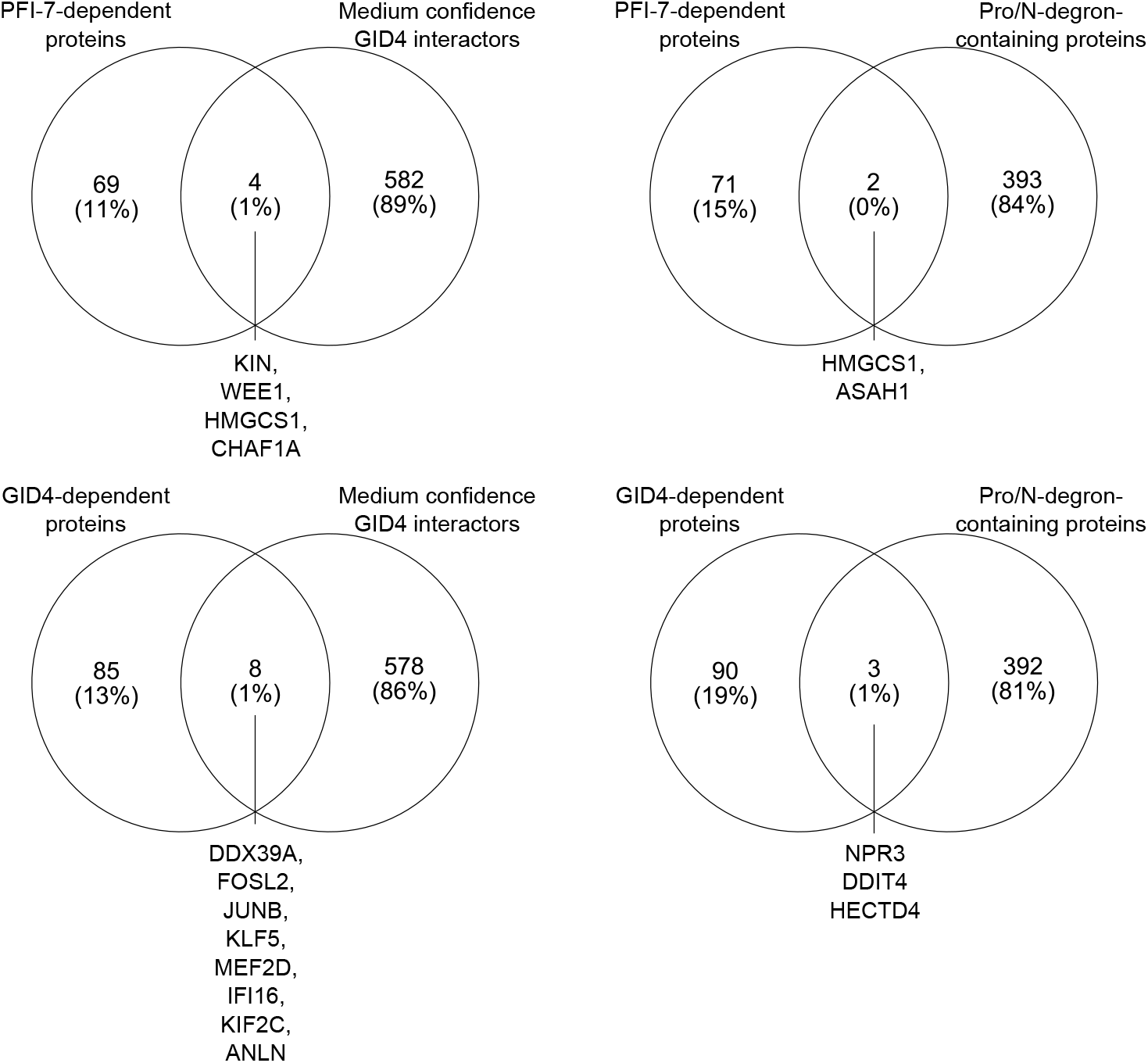
Overlap between proteins significantly changed after PFI-7 treatment or GID4 overexpression and medium confidence GID4 interactors (SP>0.6 in any condition), left. Right, PFI-7 dependent and GID4-dependent proteins overlap with Pro/N-degron-containing proteins.

